# MINREACT: an efficient algorithm for identifying minimal metabolic networks

**DOI:** 10.1101/2020.01.06.896084

**Authors:** Gayathri Sambamoorthy, Karthik Raman

## Abstract

Genome-scale metabolic models are widely constructed and studied for understanding various design principles underlying metabolism, predominantly redundancy. Metabolic networks are highly redundant and it is possible to minimise the metabolic networks into smaller networks that retain the functionality of the original network. Here, we establish a new method, MinReact that systematically removes reactions from a given network to identify minimal reactome(s). We show that our method identifies smaller minimal reactomes than existing methods and also scales well to larger metabolic networks. Notably, our method exploits known aspects of network structure and redundancy to identify multiple minimal metabolic networks. We illustrate the utility of MinReact by identifying multiple minimal networks for 74 organisms from the BiGG database. We show that these multiple minimal reactomes arise due to the presence of compensatory reactions/pathways. We further employed MinReact for a case study to identify the minimal reactomes of different organisms in both glucose and xylose minimal environments. Identification of minimal reactomes of these different organisms elucidate that they exhibit varying levels of redundancy. A comparison of the minimal reactomes on glucose and xylose illustrate that the differences in the reactions required to sustain growth on either medium. Overall, our algorithm provides a rapid and reliable way to identify minimal subsets of reactions that are essential for survival, in a systematic manner.

**Author summary:** An organism’s metabolism is routinely modelled by a metabolic network, which consists of all the enzyme-catalysed reactions that occur in the organism. These reactions are numerous, majorly due to the presence of redundant reactions that perform compensatory functions. Also, not all the reactions are functional in all environments and are unique to the environmental conditions. So, it is possible to minimise such large metabolic networks into smaller functional networks. Such minimal networks help in easier dissection of the capabilities of the network and also further our understanding of the various redundancies and other design principles occurring in these networks. Here, we have developed a new algorithm for identification of such minimal networks, that is efficient and superior to existing algorithms. We show the utility of our algorithm in identifying such minimal sets of reactions for many known metabolic networks. We have also shown a case study, using our algorithm to identify such minimal networks for different organisms in varied nutrient conditions.

## 1 Introduction

Genome-scale metabolic models (GSMMs) [1, 2] have been reconstructed for organisms from different forms of life over the last two decades [3]. GSMMs provide a wide range of insights into the metabolic capabilities of organisms, and have been exploited for a variety of applications [4–6]. GSMMs essentially comprise all known stoichiometrically balanced enzyme-catalysed metabolic reactions in an organism [1], along with other details such as the corresponding gene–protein–reaction associations. Numerous studies have been carried out to understand various organisational and design principles in metabolic networks [7–12]. Analysis of such design principles provides insights into various defining characteristics of metabolic networks, such as redundancy [10]. Metabolic networks exhibit remarkable redundancy in a wide variety of environments due to the existence of isozymes as well as alternate parallel pathways [10,13]. Besides, not all reactions in a metabolic network are functional under all conditions [14]; many reactions are active and functional only under specific nutrient conditions. Thus, it is interesting to study sets of reactions that can independently support growth under one or more conditions. To identify these sets of reactions, which can be viewed as *minimal reactomes*, a number of methods have been developed.

The earliest method, proposed by Burgard *et al* uses a simple MILP formulation that minimises the number of active reactions in the network [15]. Subsequently, graph-theoretic [16–18] and constraint-based approaches [19–21] have been developed to identify minimal metabolic networks. Some of the methods developed so far do not guarantee to identify the minimal metabolic network, rather reduce networks by removing undesired reactions. Some methods arrive at smaller networks by solving extensive MILP problems that increases the time complexity. The existing methods also do not consider known aspects of network structure and redundancy.

In this paper, we formulate a new approach to identify minimal reactomes, which leverages the network structure, particularly the redundancy between different reactions. Specifically, our approach exploits the reaction classes identified by parsimonious flux balance analysis (pFBA) [22] to prune the reactome, by removing ‘unnecessary’ reactions. We compare our approach with previous methods published and show that our approach is time efficient in the case of large networks compared to the recent method MinNW [21]. We also find that our method identifies smaller sets of reactions, *i. e.* smaller minimal reactomes, as compared to the earlier methods. Further, as an illustration of our approach, we identify multiple minimal reactomes in yeast, and illustrate the differences between the different reactomes that can support growth in the same medium. We then went ahead to identify the minimal reactomes of 70+ organisms and analysed the reactions in their minimal reactomes. Overall, our algorithm provides a new approach to efficiently and rapidly identify minimal metabolic networks from GSMMs.

## 2 Methods

### 2.1 Flux Balance Analysis

Flux balance analysis (FBA) [23,24] is a constraint-based approach for predicting the metabolic capabilities of organisms, using GSMMs. The method predicts the growth phenotypes of metabolic networks in a given nutrient condition by assuming that the network is at steady state. FBA identifies an optimal flux distribution for a metabolic network by solving a linear programming problem, maximising/minimising a given objective function, typically the growth rate of the cell. The formulation of FBA is as below:

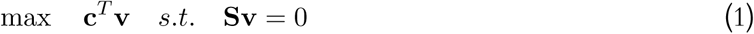

where **c** depicts the objective function, **v** is the vector of fluxes of all the reactions in the network and **S**, the stoichiometric matrix of dimensions *m* × *r*, depicts the stoichiometry of all *m* metabolites in *r* reactions present in the network.

### 2.2 pFBA

pFBA [22] is a variant of FBA. In addition to the regular FBA constraints, pFBA additionally minimises the sum of fluxes through the entire network, while also preserving the objective, such as maximal biomass production. Further, while minimising the sum of fluxes in the network, the method also classifies genes and reactions into different classes, optimising the user-defined objective. The reactions are classified as follows:

**Blocked reactions** that cannot carry a flux under the given nutrient conditions

**Zero-flux reactions** that carry only a zero flux under the given nutrient conditions

**Essential reactions** that are absolutely necessary for growth and cannot be deleted from the network

**pFBA optima reactions** that contribute to the optimal solution while minimising the flux of the reactions in the network

**Enzymatically Less Efficient (ELE) reactions** that are typically longer alternative pathways for a given metabolic conversion, and consequently have more enzymes

**Metabolically Less Efficient (MLE) reactions** that drive flux away from the cellular objective (function), thus lowering the metabolic efficiency

### 2.3 The MinReact algorithm

Every metabolic network has hundreds of essential reactions—however, just these essential reactions cannot comprise a minimal reactome. There are typically many more reactions, which are not *singly* essential, by virtue of the presence of alternative/compensatory reactions in the reactome. These reactions, therefore, comprise higher order *synthetic lethals*, such as double or triple lethals. Synthetic lethals are sets of reactions, which cannot all be simultaneously removed from the network without abolishing growth. Therefore, a minimal reactome will include all the singly essential reactions (‘single lethals’) and also multiple reactions from higher order lethals. In the minimal reactome, though, every remaining reaction must be singly lethal, as the alternative reactions have already been discarded from the (minimal) metabolic network. Reactions in a minimal metabolic network should thus ideally be chosen in a way that the total number of reactions in the network is minimised. This mathematically translates to performing a zero-norm minimisation of the flux vector, which is essentially the same as the original formulation by Burgard [15]. However, this is difficult to solve accurately, given that it is a NP-hard problem, and typically one of many possible solutions is arbitrarily obtained, on solving the problem using heuristics. These solutions are not minimal, as we will illustrate, and it is often possible to find more minimal networks.

On the other hand, our approach for identifying minimal reactomes, MinReact, exploits the reaction classes of pFBA [22], to identify minimal reactomes by exploiting network structure. A closer look at the classes of reactions identified using pFBA reveals that the MLE reactions, if present, reduce the objective flux. Obviously, the blocked reactions can be excluded from the network, as they will never carry a flux. Further, the reactions that do not carry any flux in the pFBA solution (‘zero-flux reactions’) can also be discarded from the network, without affecting network flux.

Ideally, the pFBA optima reactions constitute the essential reactions and other reactions necessary to support growth. However, due the presence of redundancies in the large metabolic networks and the existence of alternating pathways, there exist multiple groups of reactions that can yield maximal growth. Taking this into account, we framed our approach for identifying minimal metabolic networks. The approach used in MinReact is depicted in Figure 1. Our approach to identify the minimal reactome is divided into three major steps as follows:

**Figure 1:**
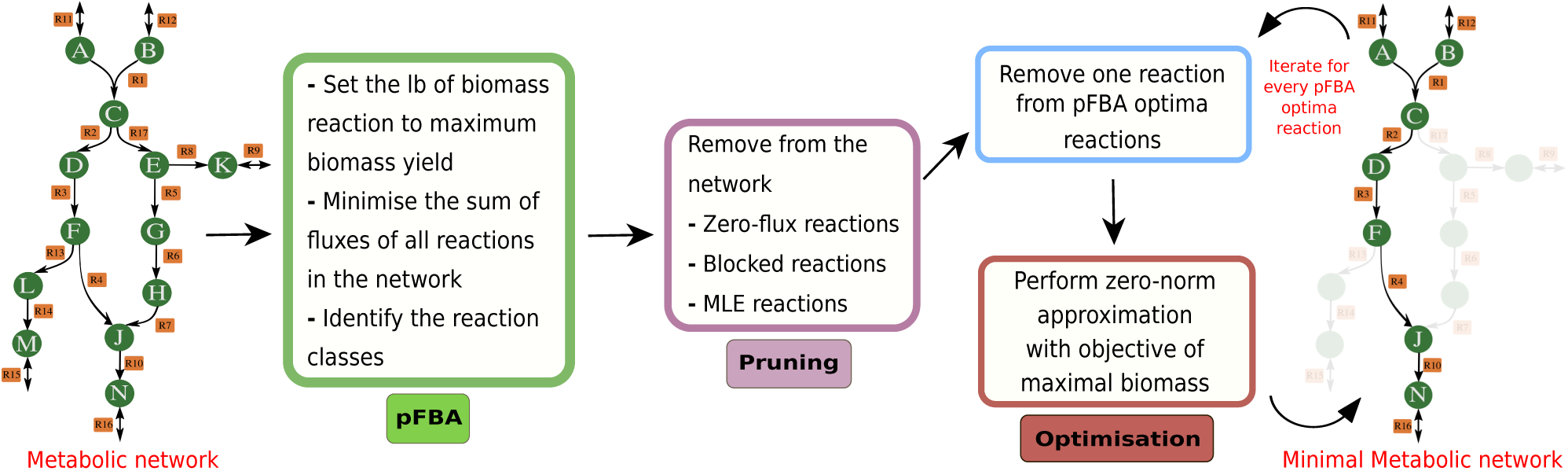
MinReact approach for identifying the minimal reactome. The left-most panel shows a sample metabolic network, where every circle represents a metabolite. The arrows represent reactions that occur, labelled with the reaction IDs; The right-most panel shows the reactions and metabolites highlighted that constitutes the minimal reactome.

1. **pFBA**: The metabolic network given as input is optimised using FBA to identify the maximal biomass growth. The sum of fluxes of all the reactions in the metabolic network is minimised while the lower bound of the biomass reaction is set to the maximal biomass growth. Further, the reaction classes of the network are identified using the optimised solution. The reaction classes thus identified by pFBA are as illustrated in §2.2.
2. **Pruning**: The blocked reactions, zero-flux reactions and MLE reactions are removed from the network. The blocked reactions and zero-flux reactions do not contribute to the network’s biomass under the given nutrient condition. MLE reactions drive flux away from the biomass; thus, removal of MLE reactions should facilitate maximal biomass.
3. **Optimisation**: The method iterates over the list of pFBA optimal reactions, deleting one reaction at a time and then performing a non-convex approximation for minimisation of the number of reactions in the network. This is done using FBA available in the COBRA toolbox v2.0 [25]. We then identify *J*_*nz*_, the set of reactions having non-zero fluxes [26] obtained in the optimised solution. This process is iterated for every pFBA optima reaction, resulting in multiple sets of *J*_*nz*_’s. These multiple *J*_*nz*_’s are the different sets of reactions that are independently sufficient for producing maximal growth under the given nutrient condition. Note that these multiple sets may vary in size (number of reactions) and also may not be distinct. Therefore, we further prune these multiple *J*_*nz*_’s to finally retain only the set/s that has/have the least number of reactions.

The algorithm also takes as input a tolerance parameter value, which is the flux threshold below which a reaction is regarded as deleted (default being 0). Another parameter is the growth-rate cut-off value that indicates the minimum percentage of the wild-type growth that the resulting minimal metabolic network should retain. The default value for the growth-rate cut-off is 100%. The method also additionally provides other features, which can be useful in different scenarios. For instance, the method can take in as input specific reactions to be retained in any given minimal reactome. This confers added advantage of preserving functionalities while identifying the set of reactions in the minimal reactome. Algorithm 1 describes the formulation of MinReact to identify minimal reactome.

### 2.4 Comparison of different methods for identifying minimal reactome

Several methods employ a constraint-based approach for identifying minimal reactomes such as, Burgard’s [15], FASTCORE [19], NetworkReducer [20] and MinNW [21]. We went ahead to compare our approach with Burgard’s, since we employ a similar technique of minimising the number of reactions in the metabolic network, and also with MinNW, as the method was proven to be better than the other methods [21]. Due to lack of implementation codes, the extended method by Burgard could not be compared.

The formulation of Burgard’s initial method [15] of minimising the number of active reactions in the network was simulated for our comparison. MinNW method [21] was performed with the codes available in literature. The parameters used for executing the MinNW method were BigM = 1, fast = 1, use_F2C2 = 1. The other sets of parameters resulted in a slower performance than the parameters considered above.

All the models available in BiGG database [27] were used for the comparison. The models from BiGG were simulated in the nutrient conditions that existed in the BiGG database. Out of the 83 organisms available in BiGG, MinReact and Burgard’s method could identify minimal reactomes for 76 organisms, while using MinNW we could identify minimal reactomes for 74 organisms. Together, we could identify the minimal reactome for 74 models by all the three methods.

All methods were implemented for models to produce atleast 99.9% of the maximal wild-type biomass growth. The tolerance level for considering a reaction to be active was set to 10^*–*7^ × wild-type biomass growth. The models were simulated preserving the ATP maintenance in the network. All the computations were performed on a workstation with Intel^®^ Core™ i7-2600 processors, CPU of 3.40GHz with 4 cores.

#### Algorithm 1 MinReact: to identify minimal metabolic networks

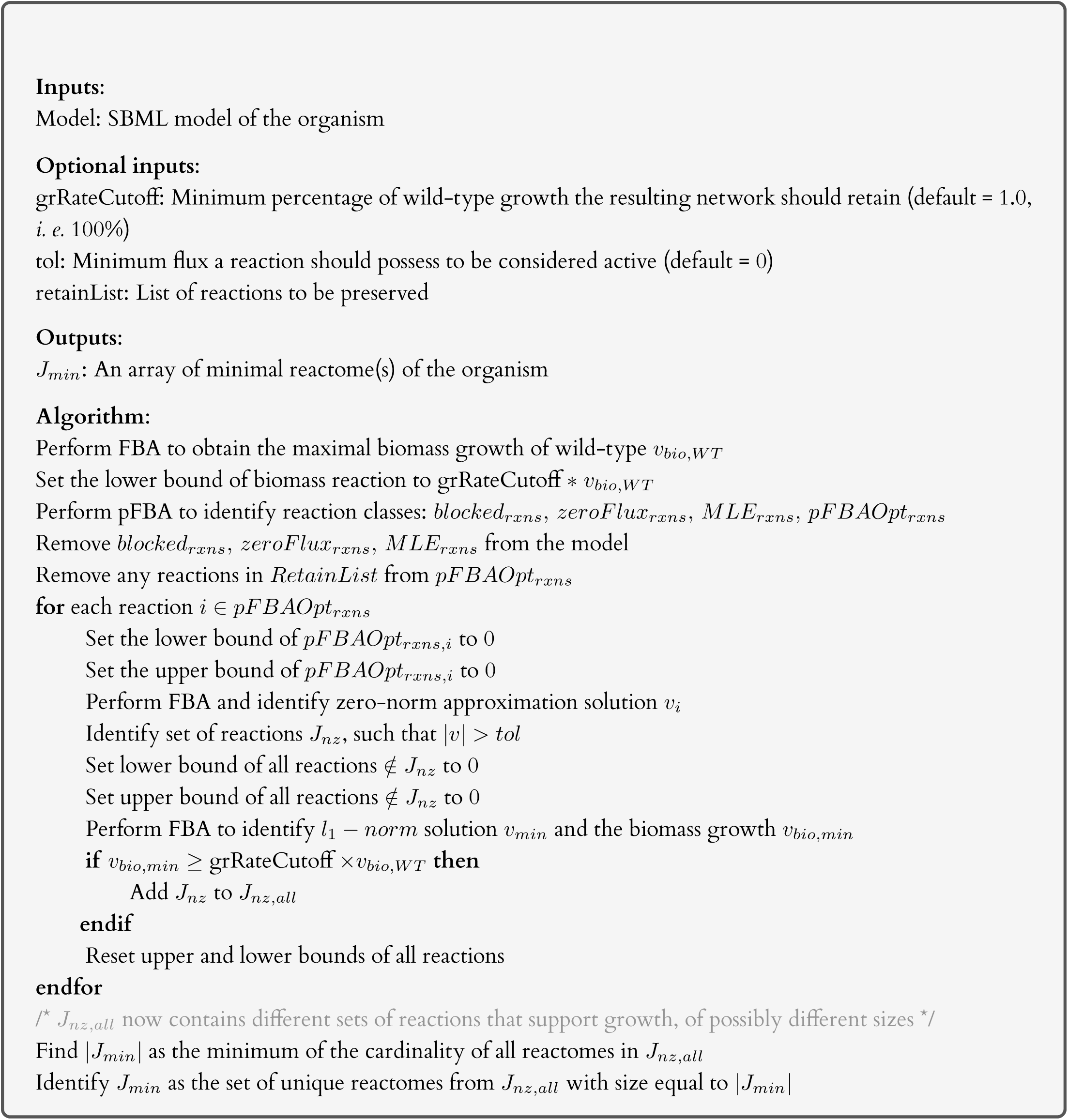

## 3 Results

In this section, we detail the performance of our MinReact algorithm, and how it compares with existing methods. Next, we show how our method identifies multiple minimal reactomes for a given metabolic model. Finally, as a case study, we identified minimal reactomes of different organisms in both glucose and xylose minimal media.

### 3.1 MinReact identifies smaller reactomes compared to other methods

We here compare the performance of MinReact with the method of Burgard [15] and MinNW [21]. Table 1 shows the number of reactions in the minimal reactome as well as the time taken to compute the minimal reactomes for different models from the BiGG database. Select models from BiGG have been reported here, with the aim to depict models from different kingdoms of life and with varied sizes of the metabolic network. The results for the remaining organisms from BiGG are detailed in Supplementary file S1.

**Table 1:** Comparison of MinReact with Burgard’s method and MinNW. The table shows the comparative performance of the three methods for different organisms. **n**_**rxn**_ denotes the total number of reactions in the metabolic network; **n**_**met**_ denotes the number of metabolites in the metabolic network. The number of reactions in the minimal reactome, |*J*_min_|, and the time taken to perform the simulations (in seconds) are tabulated for all three methods. The least number of reactions in the minimal reactome among the three methods are highlighted in bold.

| Organism | $n_{\text{rxn}}$ | $n_{\text{met}}$ | Burgard method | | MinNW | | MINREACT | |
| --- | --- | --- | --- | --- | --- | --- | --- | --- |
| | | | $\vert J_{\min}\vert $ | Time (s) | $\vert J_{\min}\vert $ | Time (s) | $\vert J_{\min}\vert $ | Time (s) |
| *Helicobacter pylori* | 554 | 485 | **313** | 0.120 | 314 | 3.85 | **313** | 13.09 |
| *Synechocystis* sp. PCC 6803 | 863 | 795 | 527 | 0.146 | **525** | 8.09 | 526 | 18.12 |
| *Mycobacterium tuberculosis* | 1025 | 825 | 416 | 0.201 | 429 | 11.80 | **414** | 20.68 |
| *Saccharomyces cerevisiae* | 1577 | 1226 | 302 | 0.269 | 304 | 49.73 | **299** | 26.53 |
| *Salmonella enterica* | 2545 | 1802 | **483** | 0.434 | 500 | 153.16 | **483** | 60.02 |
| *Shigella boydii* | 2591 | 1910 | 440 | 0.455 | 445 | 93.67 | **438** | 63.97 |
| *Escherichia coli* str. K-12 | 2712 | 1877 | 432 | 0.509 | 447 | 170.00 | **430** | 70.14 |
| *Cricetulus griseus* | 6663 | 4456 | 306 | 2.097 | 362 | 89124.63 | **264** | 613.27 |

The number of reactions in the minimal reactome identified by our method was almost always lesser than the method suggested by Burgard, although the simulation is faster than our method as shown in Table 1. For example, in *Mycobacterium tuberculosis*, our method could identify minimal reactome with 414 reactions, while that identified from Burgard’s method contains 416 reactions. In the case of *Cricetulus griseus*, MinReact could reduce the metabolic network of 6663 reactions to 264 reactions in comparison to Burgard’s, which identified 306 reactions.

In comparison with MinNW too, we found that in most cases, MinReact identified significantly smaller networks. For example, the minimal reactome of *Mycobacterium tuberculosis* found by MinNW consists of 414 reactions while MinNW identified 429 reactions. Similarly for *Cricetulus griseus*, MinNW identified 362 reactions while MinReact could find a minimal reactome of size 264 reactions. In the case of *Synechocystis* sp. PCC 6803, MinNW method identifies 525 reactions, which is lesser when compared to MinReact that identifies 526 reactions.

We compared all three methods for the smallest minimal reactome identified for all the BiGG models. Of the 74 models under study, for 61 organisms, MinReact could identify smaller minimal reactomes while MinNW identified smaller minimal reactomes for 7 organisms. For 3 organisms, MinReact and Burgard’s identified the same number of reactions in the minimal reactomes. MinReact and MinNW identified the smallest minimal reactomes for 3 other organisms.

We also calculated the difference in the minimal reactome sizes identified by the different methods. Supplementary file S1 shows that the mean difference in the size of minimal reactome between MinReact and Burgard’s method was −2.74 (ranging from 0 to −42) indicating that on an average MinReact identifies minimal reactomes with nearly three reactions lesser than Burgard’s. Similarly, we found that on comparison with MinNW, MinReact identifies on an average minimal reactomes that are 10 reactions less on average (mean difference of −10.32 with values ranging from 22 to −103).

On comparing the time taken by MinReact with MinNW, we found that for smaller models, MinNW was faster than MinReact. However, as the number of reactions in the model increases, the time taken by MinNW increases drastically. This is further depicted in the Figure 2 that compares the time taken by the methods MinNW and MinReact with the increase in the number of reactions in the metabolic network. The figure illustrates the time taken by both methods for identifying the minimal reactomes of the models in the BiGG database.

**Figure 2:**
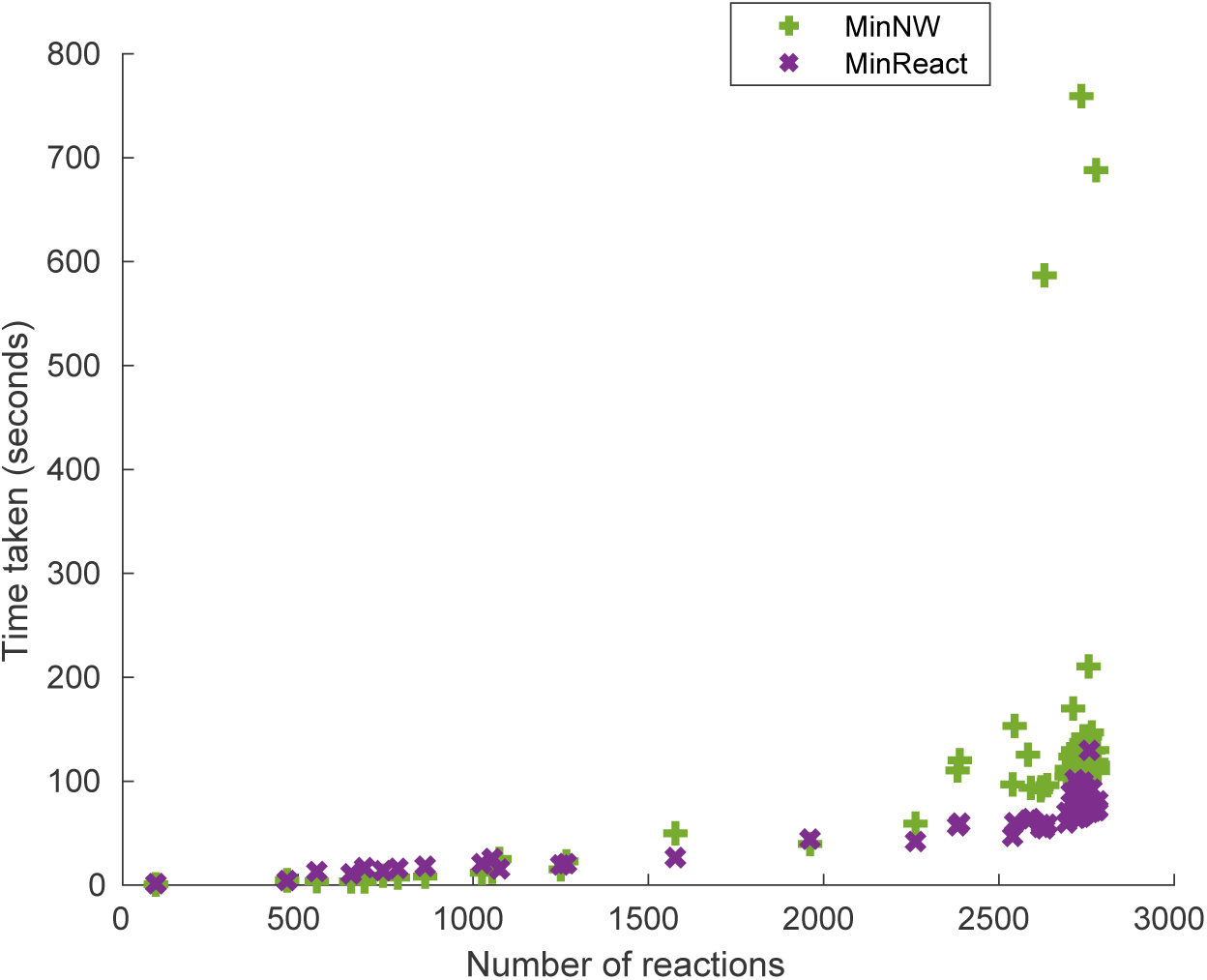
Comparison of time taken by MinReact and MinNW algorithms. The time taken to compute minimal reactomes for the BiGG models under study for both MinReact and MinNW are plotted, with increasing number of reactions in the metabolic network. One outlier point, for the largest network containing 6663 reactions, where MinNW took 89124 seconds and MinReact took 613 seconds, was excluded from the plot.

#### 3.1.1 MinReact exploits network structure to identify minimal reactomes

A closer look into the differences in the minimal reactomes by the two methods, MinReact and MinNW revealed interesting insights. As an illustration, in the minimal reactome of *Escherichia coli* str. K-12, MinNW identified 447 reactions while MinReact identified 430 reactions. On analysis of these different reactomes, we find that MinReact selects one among the compensatory reactions in such a way that the number of reactions is minimised. For example, reaction ALATA_L has a compensatory role with both the reactions VALTA and VPAMTr as illustrated in Figure 3a. Among the compensatory reactions, one reaction needs to be present for the organism to survive. These compensatory reactions are also called synthetic double lethals, since simultaneous removal of both of them causes the death of the organism [8, 13]. Similarly, triple lethal reactions consist of three reactions, whose simultaneous deletion causes cell death. We further identified the number of double lethal and triple lethal reaction sets each of these reactions occur, as found in the Figure 3a. In the above example, the reaction ALATA_L is present in the minimal reactome of MinReact while is absent in the minimal reactome identified by MinNW, thus minimises the number of reactions by one. This is achieved as pFBA predicts ALATA_L as a pFBA optima reaction. Similarly, reactions E4PD and PERD perform compensatory roles with both 4HTHRK and 4HTHRA, thus forming synthetic double lethals as depicted in Figure 3b. The minimal reactome identified by MinReact consists of reactions E4PD and PERD and we find that they are a part of many triple lethals (as shown in the figure).

**Figure 3:**
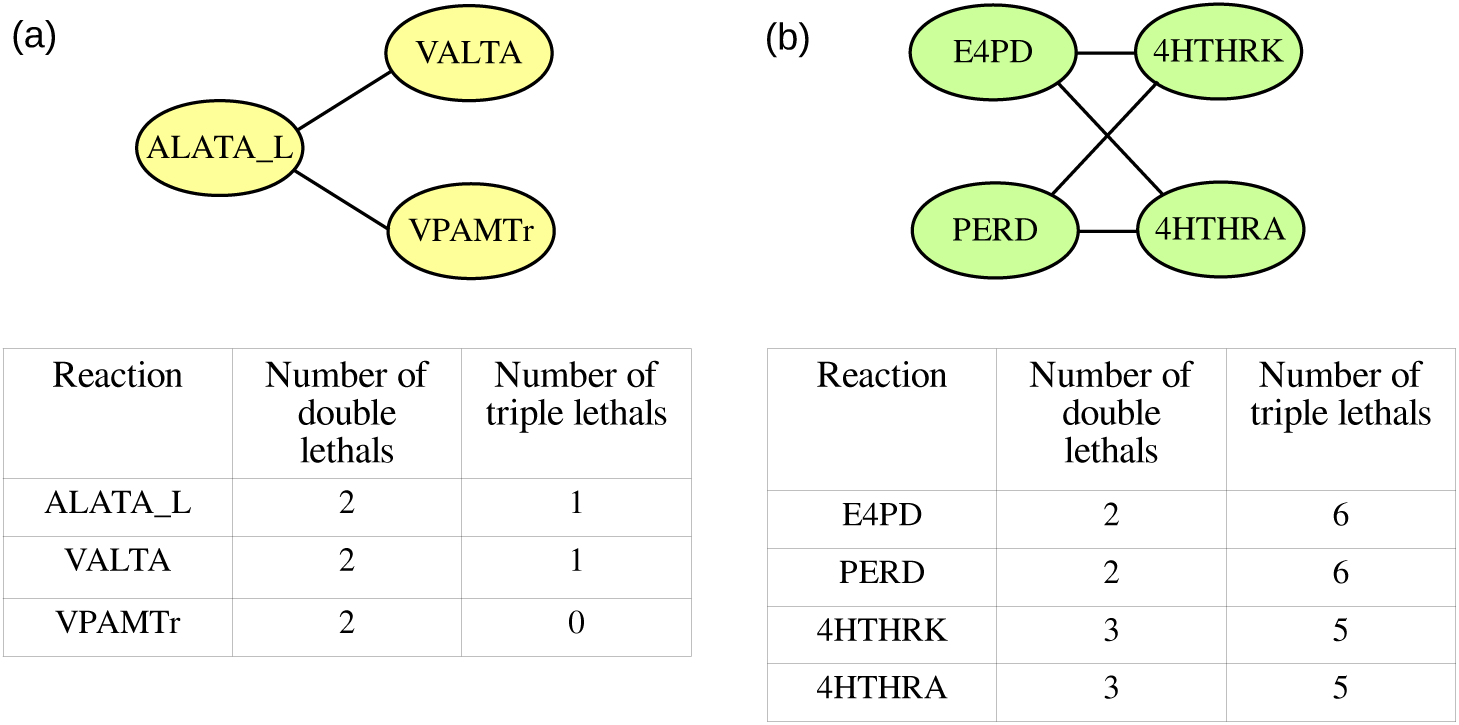
Lethal reactions in *E. coli*. The figure represents select compensatory reactions in *E. coli* iML1515 GSMM. The ovals represent the reactions and straight lines connect reactions if they exist as a double lethal. The table below illustrates the number of double lethals and triple lethals that the reaction occurs in. Panels (a) and (b) show different examples of compensatory reactions in the organism.

In any given minimal reactome, every reaction must be essential for maximal growth. That is, the removal of any of the reactions from a given minimal reactome, should result in a reduction in (or even loss of) growth. To study this, we performed single reaction deletions from every minimal metabolic network identified by each of the three methods, and report these numbers in Supplementary File S1. We found that for almost all organisms, every reaction present in the minimal reactomes identified by MinReact and Burgard’s are essential. However, not all the reactions present in the MinNW minimal reactomes were found to be essential. This illustrates that the minimal reactomes identified by MinNW are not truly *minimal*, and further reactions can be removed to identify a smaller minimal reactome. Thus, the above explains that MinReact surpasses all other methods in terms of systematically choosing the reactions that should occur in the minimal reactome.

### 3.2 MinReact can identify multiple minimal reactomes for a given metabolic network

Our approach used in MinReact is iterative and thus enables us to identify multiple reactomes for a given metabolic network. These minimal reactomes have the same number of reactions but the reactions themselves will vary. Figure 4 illustrates the number of distinct minimal reactomes that could be identified in the 77 BiGG models using MinReact.

**Figure 4:**
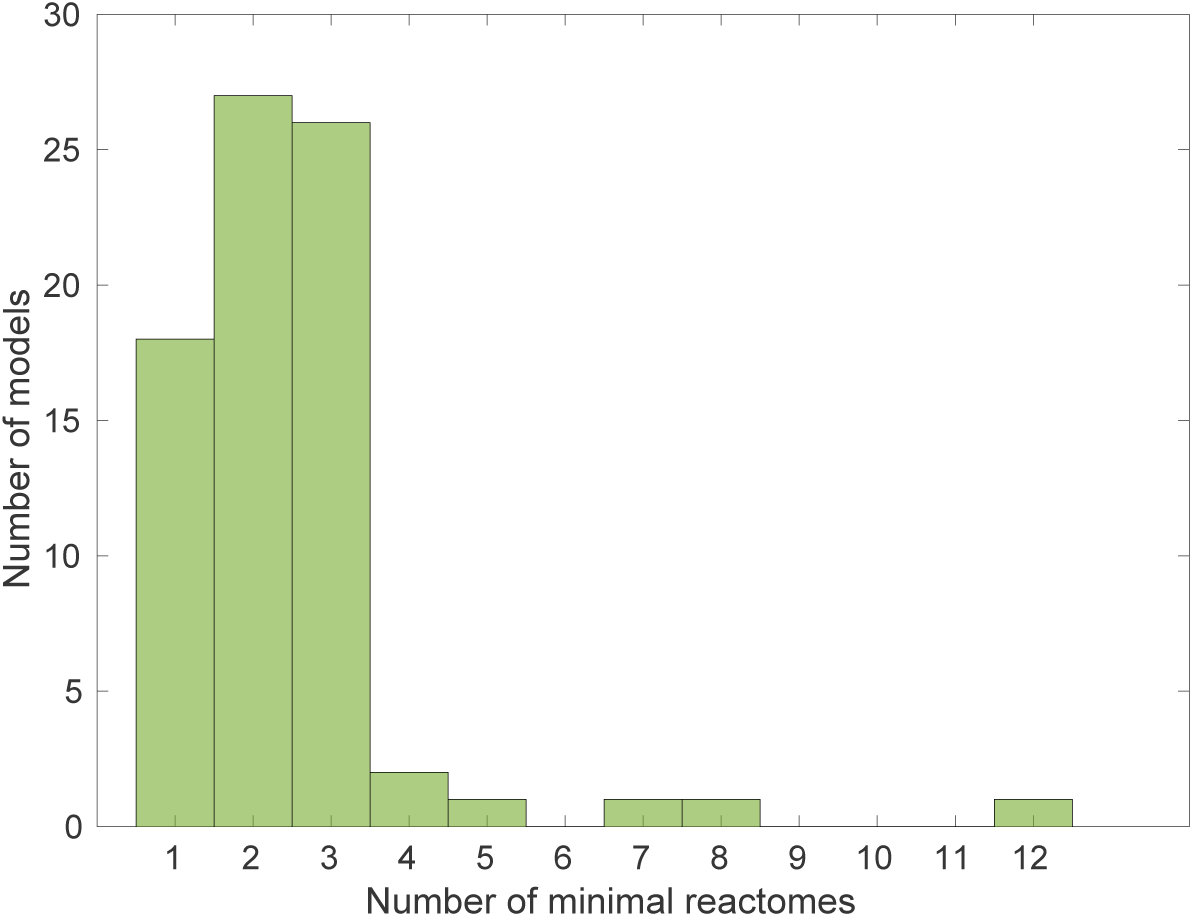
Number of minimal reactomes identified using MinReact. The figure illustrates the number of minimal reactomes that could be identified by MinReact in different models in BiGG database.

Of the 77 organisms, we could identify more than one minimal reactome for 59 organisms. For as many as 27 organisms, MinReact identified two minimal reactomes and 26 organisms had three minimal reactomes. We could identify 7 minimal reactomes for *Escherichia coli* ED1a, 8 minimal reactomes for *Salmonella enterica* and 12 different minimal reactomes for *Saccharomyces cerevisiae* S288C (iND750). As an illustration, we analysed the 12 minimal reactomes of *Saccharomyces cerevisiae* S288C iND750 model that constitutes of 278 reactions each. The metabolic network consists of 182 essential reactions, that the network cannot bypass, to grow in the nutrient condition composed of glucose, water, ammonia, oxygen, phosphate and sulphate. First, we find that across the 12 minimal reactomes, 266 reactions are common. That is, all the 12 minimal reactomes contain this set of 266 reactions. These 266 reactions comprise of the 182 reactions that are *highly* essential, and in addition contain other reactions necessary for supporting growth. On the other hand, the remaining 12 reactions (278 – 266) differ from one reactome to another.

We identified how these 12 minimal reactomes, each of size 278, vary by analysing the reactions present in them. We found the number of reactions that are common among all the pairs of 12 minimal reactomes. Of the 66 pairs, majority differ in three reactions. The least number of reactions shared between any two minimal networks was 272. We investigated the difference between two reactomes that varied maximally, with a variation in 6 reactions. One major difference was the two different ways 1-acyl-sn-glycerol 3-phosphate was formed. This metabolite can be produced from dihydroxyacetone phosphate in two steps via two alternating pathways. Either dihydroxyacetone phosphate is coverted to glyceraldehyde-3-phosphate and then to 1-acyl-sn-glycerol 3-phosphate or dihydroxyacetone phosphate can be converted to 1-acyl-glycerone 3-phosphate, that can produce 1-acyl-sn-glycerol 3-phosphate.

Further, a maximum of 277 reactions were shared between very similar minimal reactomes, thus vary in only one reaction. A closer look into the reaction that was different between minimal reactomes pertained to double or higher order lethals in the organism. For example, reactions ADK3 and NDPK1 form double lethal in the organism and they are found in two different minimal reactomes that differ in only one reaction. Reactions G5SD and G5SD2 are a part of a triple lethal in the network and are found in different minimal reactomes of equal length. Similarly, we found that {AADSAD1, AADSAD2}, {PPND,PPND2} are double lethals present in different minimal reactomes differing in only one reaction.

### 3.3 Minimal reactomes of different organisms in glucose and xlyose

How different is the metabolic core of different organisms in the same nutrient condition? How different are minimal reactomes that occur in different environments? Can we identify reactomes that are most similar between two nutrient conditions? To address these questions, we identified the minimal reactomes of different organisms in glucose and xylose environment. Table 2 details the minimal reactome size using MinReact for different organisms considered in both glucose and xylose minimal medium. The minimal reactome size varied for the different organisms and the percentage of reactions in the minimal reactome out of the total reactions present in them varied between 12–27% in both glucose and xylose. An interesting finding is that *Bifidobacterium longum* had the smallest minimal reactome compared to all the other organisms in both glucose and xylose. It is interesting that different organisms possess different numbers of minimal reactomes in the same nutrient conditions. This portrays the fact that organisms exhibit varying levels of redundancy, particularly for their bare minimum functions. *Salmonella enterica* had the maximum number of minimal reactomes in both glucose and xylose.

**Table 2:** Comparison of minimal reactomes of different organisms on glucose and xylose. |***J***|, total number of reactions in the metabolic network; |***J***_***min***_|, the number of non-zero reactions after optimisation—corresponds to the number of reactions in the minimal reactome; ***n***_**min**_, number of minimal reactomes identified (of size |***J***_***min***_|). The number of reactions shared between all pairs of minimal reactomes identified on glucose and xylose were found, and their range is shown in the last column.

| Organism | $\vert J\vert $ | On glucose | | On xylose | | Number of reactions shared |
| --- | --- | --- | --- | --- | --- | --- |
| | | $\vert J_{min}\vert $ | $n_{min}$ | $\vert J_{min}\vert $ | $n_{min}$ | |
| *Escherichia coli* str. K-12 | 2583 | 438 | 2 | 438 | 1 | 429–430 |
| *Saccharomyces cerevisiae* | 1577 | 297 | 2 | 299 | 3 | 282–286 |
| *Klebsiella pneumoniae* | 2262 | 313 | 1 | 315 | 1 | 306 |
| *Bacillus subtilis* | 1250 | 333 | 2 | 335 | 2 | 327–331 |
| *Salmonella enterica* | 2545 | 483 | 8 | 485 | 7 | 477–480 |
| *Bacteroides eggerthii* | 2069 | 264 | 1 | 264 | 3 | 250–251 |
| *Bifidobacterium longum* | 1005 | 257 | 1 | 258 | 3 | 250–251 |
| *Citrobacter amalonaticus* | 1827 | 495 | 8 | 498 | 1 | 485–489 |
| *Enterobacter aerogenes* | 1840 | 500 | 3 | 502 | 1 | 489–492 |
| *Parabacteroides merdae* | 2474 | 328 | 2 | 330 | 3 | 314–321 |

We observed that there were 78 reactions common between all 10 minimal reactomes of different organisms in glucose. In the xylose medium, the number of reactions common across different organisms was 83. We found that reactions belonging to the phenylalanine metabolism and nucleotide interconversion were predominant among the pathways of the common reactions.

We went ahead to analyse the multiple minimal reactomes of *Salmonella enterica* in glucose and xylose environments. In glucose, the number of minimal reactomes identified were 8, each with 483 reactions, while in xylose, the number of minimal reactomes identified were 7, each with 485 reactions. We analysed the similarity of the minimal reactomes in glucose and xylose by studying the reactions that occur in the different reactomes. We found that out of the 56 combinations, the maximum number of reactions shared between minimal reactomes of glucose and xylose was 480. The minimum number of reactions shared between all combinations of minimal reactomes in glucose and xylose is 477 reactions.

We investigated the minimal reactomes that are maximally similar and found that they differ only in the reactions specific to their nutrient conditions, namely glucose and xylose. Reactions EX_glc_D_e, GLCptspp and GLCtex that are specific to conversion of glucose were present in the minimal reactome of glucose and absent in xylose. Similarly, reactions specific to xylose uptake, namely, EX_xyl_D_e, XYLI1, XYLK, XYLtex, XYLt2pp were present only in the minimal reactome of xylose. Hence, while minimising the number of reactions in the metabolic network, the core reactions necessary for the survival are retained and only nutrient specific reactions differ between minimal reactomes in different nutrient conditions. Thus, using the above illustration, we could find minimal reactomes in different nutrient conditions and also identify those that vary minimally in the two nutrient conditions.

## 4 Discussion

Genome-scale metabolic networks have been studied to obtain various insights into the organisation of metabolic networks of diverse organisms. Many previous studies have uncovered interesting design principles of these networks, notably redundancy [8, 10, 12]. Given this redundancy, it is easy to imagine that metabolic networks contain far more reactions than are expressly necessary for growth in any given environment. Thus, it is of interest to identify minimal sets of reactions, or minimal metabolic networks, which can support growth in any given environment.

A few constraint-based methods have been developed in the past to construct minimal metabolic networks from GSMMs [19–21]. The optimisation problems formulated for this purpose are difficult to solve, and notably have more than one minimum. Most algorithms identify only a single (arbitrary) minimal reactome for a given GSMM. In this study, we develop a simple method, that builds on the widely used parsimonious formulation for FBA, to identify multiple minimal reactomes in a given GSMM, in a systematic fashion. Our approach, MinReact identifies the minimal metabolic networks within a given GSMM, for the production of biomass components, by eliminating unnecessary reactions delineated by pFBA, and assembles multiple minimal reactomes.

We show that MinReact typically identifies minimal metabolic networks of smaller sizes than existing algorithms. Further, we show that MinReact scales better to larger metabolic networks, compared to existing methods such as MinNW. Although the early method of Burgard [15] is faster than MinReact, our method often produces superior networks, identifying the most minimal pathways that should comprise a given GSMM. Importantly, we have a principled way of identifying minimal metabolic networks which exploits network structure and redundancy, identifying smaller networks. Notably, our iterative approach identifies multiple possible minimal reactomes of the same size, for a given GSMM.

Minimal reactomes identified for varied organisms on the same nutrient condition are different in terms of the size and also on the number of minimal reactomes. This shows that organisms possess different ways of metabolising nutrients and also the varied levels of redundancy that exist. Further, analysing the minimal reactomes of an organism in different nutrient conditions can aid in identifying reaction sets that are minimal and yet can support growth in those nutrient conditions. This paves way to identify minimal set of reactions that can satisfy multiple functionalities. This is useful, say for example, in metabolic engineering applications, if we intend to identify a minimal set of reactions that can grow on two different nutrient conditions.

Minimal metabolic networks help in identifying the core set of reactions that organisms should retain in a given nutrient condition. Complex analysis of large-scale networks, such as identification of EFMs also becomes easier with such minimal networks. Exploring the metabolic network space for interventions in metabolic engineering applications becomes easier with a smaller set of reactions preserving the desired functionality. Overall, our method MinReact identifies metabolic networks efficiently and in a systematic manner. The study of such minimal metabolic networks gives various insights into the versatility of metabolic networks, and also furthers our understanding of the underlying design principles.

## Supplementary Information

### Supplementary file S1. Minimal reactomes of 83 organisms in the BiGG database

The number of reactions in minimal reactome for all the BiGG organisms identified by the three different methods, Burgard’s, MinNW and MinReact have been documented. The time taken by each of the method is also documented. The number of essential reactions in the minimal reactomes identified are tabulated.

## Acknowledgements

G.S. acknowledges the Initiative for Biological Systems Engineering (IBSE), IIT Madras, India for the PhD Studentship.

